# Genome biology of long non-coding RNAs in humans: a virtual karyotype

**DOI:** 10.1101/2024.09.23.613004

**Authors:** Alessandro Palma, Giulia Buonaiuto, Monica Ballarino, Pietro Laneve

## Abstract

**Background:** Long non-coding RNAs (lncRNAs) represent a unique and groundbreaking class of RNA molecules that exert regulatory functions with remarkable tissue and cellular specificities. Although the number of identified functional lncRNAs is increasing, comprehensive profiling of lncRNA genomics remains elusive. Creating a virtual lncRNA karyotype is especially important for species whose intrinsic features enable their biosynthesis and function in context-dependent manners.

**Results and conclusions:** To address this challenge, we employed existing annotation files to create a statistical genomics portrait of lncRNA genes for comparison with protein-coding genes. We provide a foundational reference for exploring the non-coding genome, offering insights into the genomic characteristics of lncRNAs that may enhance understanding of their biological significance and impact.

## Background

Long non-coding RNAs (lncRNAs) emerged in the early to mid-2000s as surprising evidence of the genomes’ transcriptional pervasiveness. Initially considered junk material, their importance as precise spatiotemporal tuners of gene expression has become increasingly recognized with the accumulation of functional data, reshaping our mechanistic interpretation of the genetic information flow (Mattick et al., 2023). The lncRNA attributes, including their high expression specificity and ability to scaffold chromatin, RNA, and proteins underlie their influence on cellular identity and activity (Statello et al., 2021; Yao et al., 2019).

Additionally, their propensity for rapid evolution contributes to biological complexity. The extent of this potential underscores the necessity of detailed annotations to provide essential context and information for the functional descriptions of lncRNA genes (lncG), as well as their genetic variations that can impact health and disease. However, lncRNAs remain relatively understudied from this standpoint (Amaral et al., 2023). In particular, the lack of comprehensive structural and functional annotations hinders the investigation of their roles in cellular processes and functional enrichment studies derived from increasingly widespread next-generation sequencing data.

As the primary structure of lncRNAs shapes their biological properties and binding to specific partners, many computational tools have been developed for its analysis (as reviewed in (Ballarino et al., 2023)). Notably, sequence features in introns retained after splicing completion are of increasing interest, as these have been shown to influence the scaffolding activities and the specific binding of lncRNAs to macromolecular interactors (Desideri et al., 2020; Dey and Mattick, 2021; Dumbović et al., 2021; Ribeiro et al., 2018; Zhen et al., 2023).

LncRNA classification was initially based on the lack of coding potential (Laurent et al., 2015), although recent discoveries have revealed that certain lncRNAs can encode functional micro-peptides (reviewed in (Anderson et al., 2015; Hartford and Lal, 2020; Pan et al., 2022)). Furthermore, a recent consensus proposed classifying these transcripts as being longer than 500 nt, to exclude a range of small RNAs (e.g., RNA polymerase III transcripts, snRNAs, and intron-derived snoRNAs) that are 50-500 nucleotides in length (Mattick et al., 2023). Overall, it emerges that the classification of lncRNAs is not always clear-cut due to their functional heterogeneity, necessitating a deeper study of their genomic organization to gain a more comprehensive understanding of this RNA class.

In the human genome, protein-coding genes (PCGs) and lncGs are distributed along nearly 3.2 billion base pairs of DNA, arranged across the 46 chromosomes in a precise manner. The genomic arrangement of lncGs has not been extensively studied yet, but it is plausible that it could affect the role of lncRNAs as regulators of gene expression. To address this issue, we examined the genomic organization of lncGs in the human genome using annotation data from genomic databases including GENCODE (Frankish et al., 2023), Ensembl (Martin et al., 2023) and dbSNP (Sherry et al., 2001), and captured genomic features like chromosomal distribution, nucleotide content, and point mutations with possible roles in lncRNA biology.

## Results

### Distribution of lncGs along the genome

To provide possible insights aiding the study of functional significance of lncGs, we first analyzed their distribution across the human genome. In line with the FANTOM5 project (Abugessaisa et al., 2021), the current GENCODE catalogue (version 46) lists a comparable number of human long non-coding (20,310) and protein-coding (20,065) genes, although with different coverage on the genome (614,682,854 nt for lncGs, and 1,380,521,881 for PCGs). Despite their positive correlation (R=0.68), their distribution among chromosomes is non-uniform (**Supplementary Figure 1a**). Specifically, chromosomes 19, 11, and X exhibit significantly more PCGs, whereas chromosomes 2, 5, and 8 display a higher number of lncGs (**Figure 1a**). Chromosome 1 harbors the highest absolute number of lncGs (1,649), followed by chromosome 2 (1,451), chromosome 12 (1,150), and chromosome 5 (1,118) (**Supplementary Figure 1a** and **Supplementary Table 1**). In contrast, the Y chromosome contains the fewest lncGs (112), which are restricted to its short arm and to a proximal region of the long arm. Additionally, chromosome length also appears to be correlated with lncG abundance (R=0.58), albeit with some exceptions. For instance, chromosome X contains fewer lncGs than expected based on its large size (**Figure 1b**), while the smaller metacentric and submetacentric chromosomes 16, 17, and 19 exhibit a higher number of lncGs relative to their length. Notably, chromosome 16 stands out as the second most lncG-dense chromosome, with approximately 10.5 lncGs per megabase (Mb).

**Figure 1.**
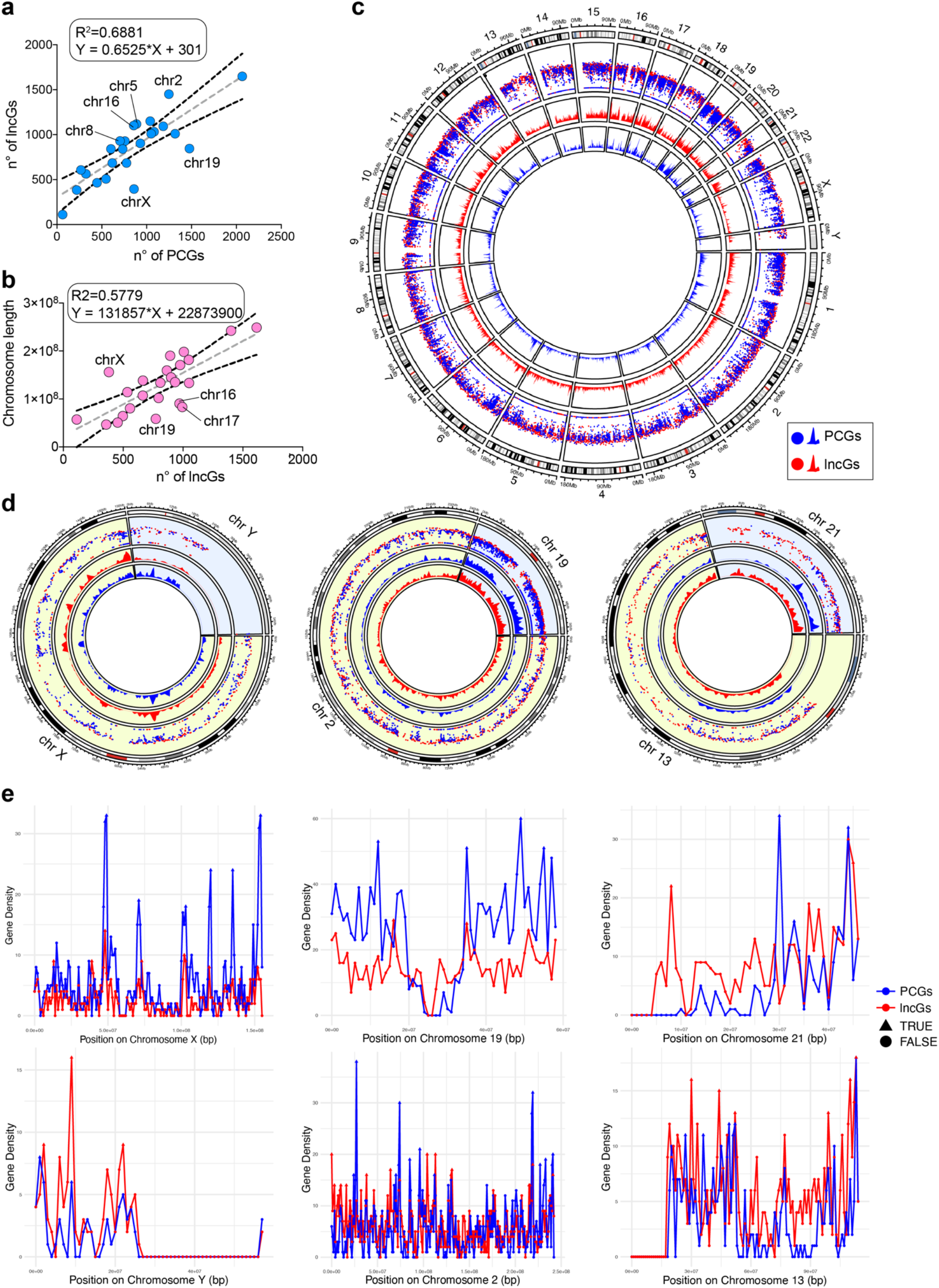
Chromosomal distribution of protein-coding and long non-coding RNA genes. Correlation between the number of long non-coding RNA (lncGs) and **(a)** protein-coding genes (PCGs) or **(b)** chromosome length. **(c)** Rainfall plot showing the distribution of human protein-coding (PCG; blue) and long non-coding RNA (lncG; red) genes across chromosomes. **(d)** insets on chromosomes X, Y, 2, 19, 13 and 21 depicted below. Dots represent individual genes, while peaks represent distributions within a 1 Megabase window. **(e)** Peaks quantification of the density of lncGs and PCGs in the corresponding (insets) chromosomes (see **Methods** for the quantification methodology).

Furthermore, an analysis of the overall lncG densities (calculated as the number of lncGs per 1 Mb for each chromosome), revealed high densities and local peaks of lncGs or PCGs in certain chromosomal regions (**Figure 1c, d**).

**Supplementary Figure 1.**
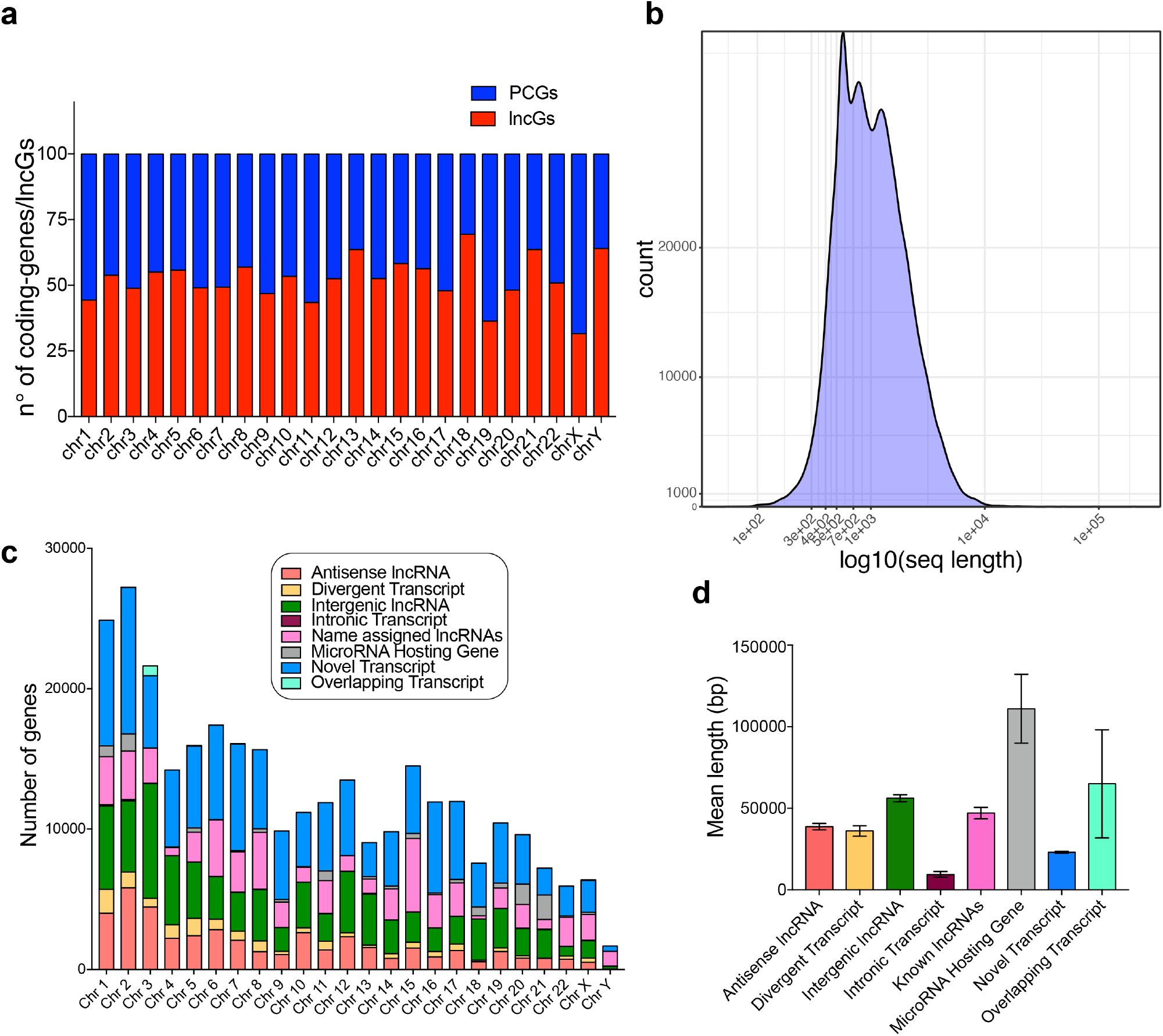
**(a)** Stacked barplot showing the percentages of coding and long non-coding RNA genes across chromosomes. **(b)** Distribution of lncRNA length showing the existence of three distinct peaks centered around 600, 800 and 1500 base-pairs. **(c)** Stacked barplot reporting the number of genes belonging to each lncRNA subtype across the different chromosomes. **(d)** Mean length of lncGs grouped by subtype.

Notable examples of lncG local peaks are evident on chromosome 18. Conversely, a higher density of PCGs with marked local peaks is found on chromosomes 13 and X (**Figure 1c, d**). Other chromosomes show comparable densities between coding and long non-coding RNA genes. For instance, chromosome 19 exhibits a lncGs density (845, corresponding to 14.42 per Megabase (Mb)) that partially reflects that of coding genes. PCGs are extremely numerous on chromosome 19 as well (1,477 genes, 25.2 genes/Mb) (**Figure 1a**), in line with chromosome 19 being considered the most gene-dense human one (Grimwood et al., 2004). A unique case is represented by chromosome 21, which exhibits a distinct separation between a region enriched in lncGs, on its short arm and the proximal part of the long arm, and a region abundant in PCGs, the latter situated in the distal part of its long arm (**Figure 1c, d**). Another key feature is present on Y chromosome, which displays a higher lncGs’ density compared to PCGs.

Collectively, these findings indicate that lncGs are almost unevenly distributed throughout the human genome, with specific chromosomes or chromosomal regions that appear enriched in either lncGs or PCGs. The presence of local peaks of lncGs along chromosomes hints at positional clustering which could potentially impact lncRNA expression and function.

### Long non-coding genes: about size and nucleotide content

To further delve into more sequence-specific features, we analyzed the lengths of PCGs and lncGs. Our findings revealed a significant variability in lncG length, averaging 31,634 nucleotides (nt), which was quite dependent on chromosome size, and tended to be shorter than PCGs averaging 72,443 nt in length (**Figure 2a**). Shorter lncGs were typically located on smaller chromosomes such as 16, 17, 19, and 22, while longer lncGs were found on larger ones such as chromosomes 4 and X (**Figure 2a**), with few exceptions. For instance, despite chromosome Y being the third smallest human chromosome, it contains 112 lncGs (mean length: 31,439 nt). Their length is comparable to the average length of lncGs across all chromosomes. Conversely, chromosome 1, the largest human chromosome spanning 248,956,422 nt, harbors the highest number of lncGs, totaling 1,649. However, the average length of these lncGs is 26,902.7 nt. This value falls significantly below the overall mean length of lncG from all chromosomes.

**Figure 2.**
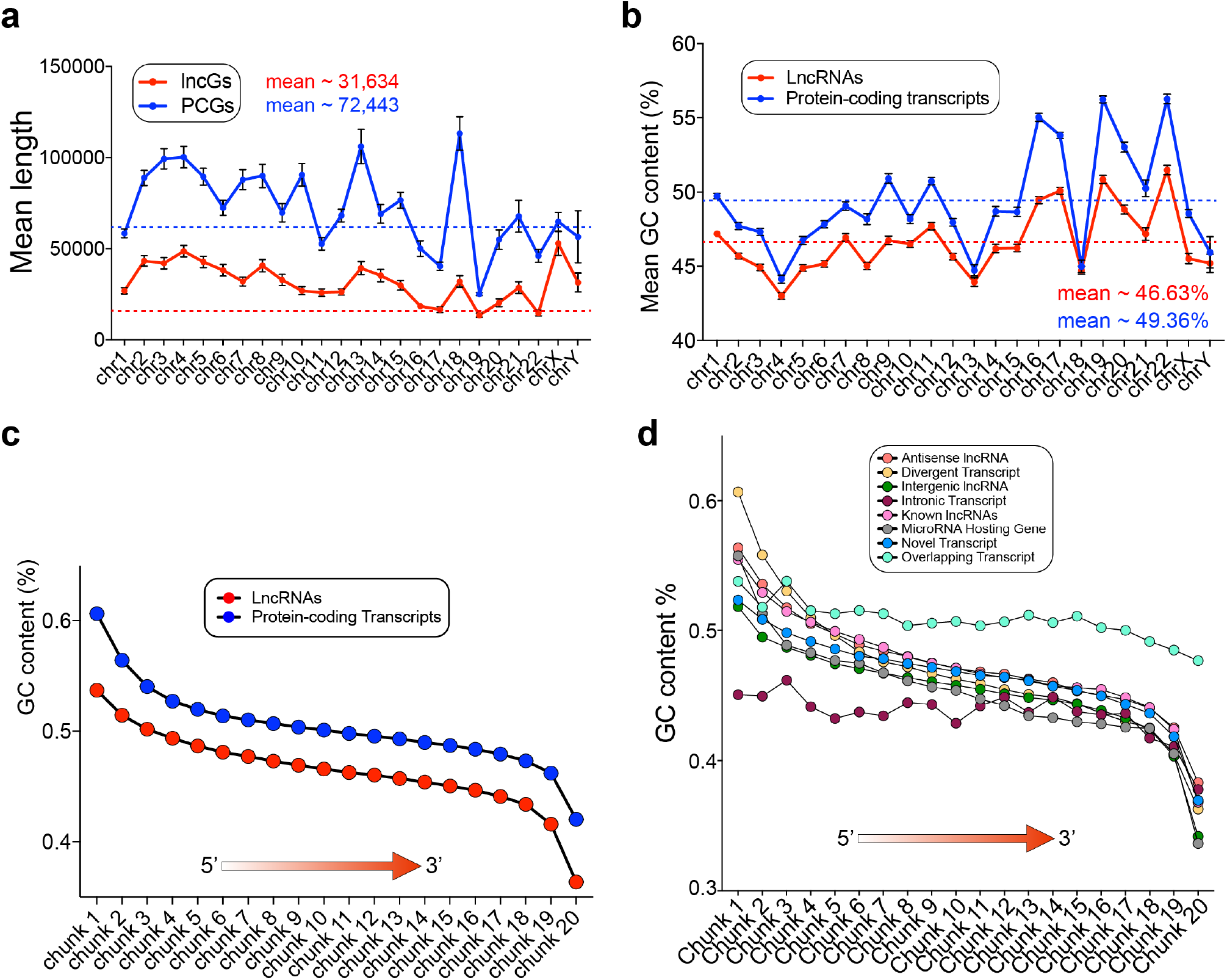
Length and GC content of long non-coding and protein-coding transcripts. **(a**) Average length and(b)GC content of long non-coding and protein-coding transcripts across chromosomes. **(c)** GC skew of coding and long non-coding transcripts divided by 20 chunks proportional to each transcript length from 5’ to 3’. **(d)** GC skew of long non-coding transcripts for the distinct lncRNA subtypes.

As for the transcript sequences, we examined all the long non-coding transcripts that are annotated on GENCODE and observed that lncRNAs exhibited substantial heterogeneity, boasting a mean length of 1,319 nt and a median length of 965 nt. Notably, the length distribution of lncRNAs displays three discernible peaks (**Supplementary Figure 1b**), with the first peak around 600 nt, the second peak around 800 nt, and a third peak at 1,500 nt. This distribution hints at the possibility for classifying these transcripts into three different subcategories, based on their length. Moreover, it is worth noting that most of lncG sequences contain introns.

We then zoomed in the nucleotide composition. Regions of DNA characterized by high guanine and cytosine contents (GC-rich regions) are recognized for their significance in the formation of stable structures, thereby influencing histone deposition and genome functionality (Zhou et al., 2011). These regions often exhibit a high density of CpG dinucleotides, serving as regulatory islands for a majority of human genes (Illingworth and Bird, 2009). GC skew is frequently linked to unmethylated human CpG island promoters (Ginno et al., 20125), and their transcription can instigate the formation of R-loops stabilizing the nascent RNA at its DNA locus and generating GC-rich RNA:DNA hybrids (Hartono et al., 2015). The presence of these structures potentially underlie the role of lncRNAs as epigenetic regulators, as already shown for selected species (Ariel et al., 2020; Cloutier et al., 2016; Gibbons et al., 2018; Niehrs and Luke, 2020).

Earlier analyses have indicated that lncRNAs typically show lower GC content compared to protein-coding transcripts, with exons generally bearing a higher GC content than introns (Haerty and Ponting, 2015). However, this distinction should be interpreted cautiously and scrutinized for potential analytical biases, as some lncRNAs retain introns in their functional form (Desideri et al., 2020). The average GC content of the human genome is approximately 41% (Cohen et al., 2005; Lander et al., 2001; Piovesan et al., 2019), with chromosomes 19 and 22 characterized by the highest percentage (47.94% and 47%, respectively). When assessing the GC content of long non-coding transcripts and PCG sequences across different chromosomes, we noted an overall mean of 46.6% for lncRNAs, reaching values as high as 49.4% for protein-coding transcripts (**Figure 2b**). When looking at lncRNAs individually in each chromosome, chromosomes 16, 17, 19, 20 and 22 stood out with the highest GC content, surpassing the 50% threshold, while chromosomes 4 and 13 displayed relatively lower GC content at 43% and 43.9%, respectively. To investigate whether this GC content was maintained in the fully spliced form of lncRNAs, we examined the positional effect of GC skew. Employing a 10 nt sliding window, we found that GC content was much higher near the 5’ end of the sequence for both coding and non-coding transcripts, with the former bearing a higher GC content. In contrast, GC content significantly decreased toward the 3’ end, reaching values as low as 36.3% for lncRNAs (**Figure 2c**). As for protein-coding genes, the highest content of GC peaks at the 5’ end may be associated either with the transcriptional regulation or the nuclear export of lncRNAs. The study of the evolutionary origin, maintenance and distribution of these regions across lncRNA genes is crucial and will help to understand alterations in lncRNA expression in pathological contexts. Interestingly, we noted a tendency for GC content to be lower in nuclear-localized lncRNAs when compared to cytoplasmic ones (nuclear lncRNAs mean=49.1%, cytoplasmic lncRNAs mean=50. 4%, **Supplementary figure 2a**). However, the subcellular localization of these lncRNAs seems to be generally independent to GC content (ANOVA test adjusted p-value=0.15).

**Supplementary Figure 2.**
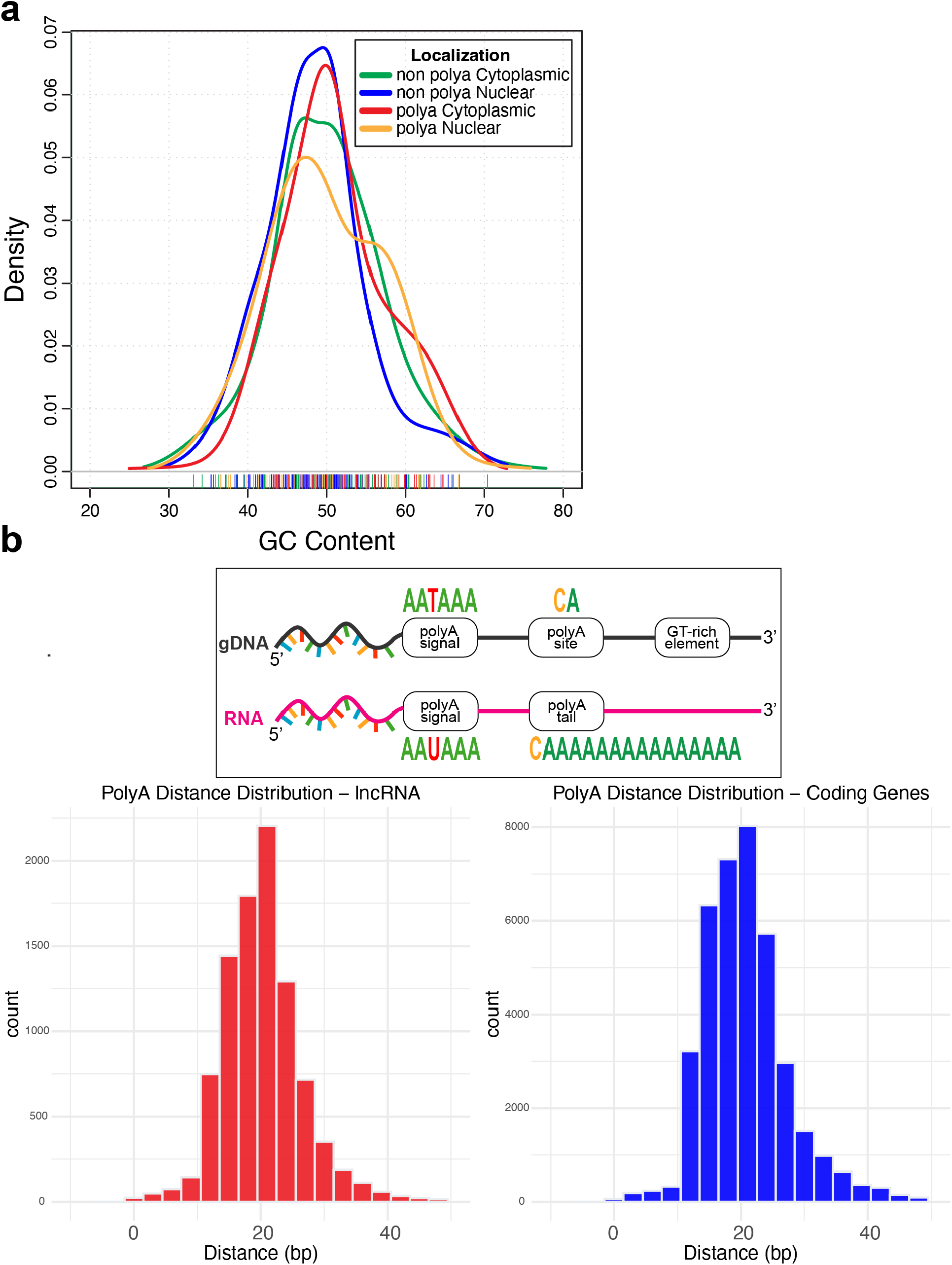
**(a)** Distribution of the GC content for polyadenylated and non-polyadenylated, cytoplasmic and nuclear long non-coding transcripts. **(b)** Distribution of the distance between polyA sites and signals for long non-coding (left) and coding (right) genes.

## Functional classification of lncG types

One significant classification of lncGs hinges on their proximity to nearby PCGs (Laurent et al., 2015). In an effort to refine the suggested nomenclature for naming lncGs (Seal et al., 2020), we analyzed their genomic features. Here, we introduce the designation “Novel transcripts” for lncGs not fitting into any of the suggested classes and identified solely by an Ensembl ID as the annotated gene name. On the other hand, “Name assigned lncRNAs” were attributed to those with an annotated gene name and a documented functional characterization (**Table 1**).

**Table 1.** LncG classification. LncG subtypes are reported together with a short description, the prefix or suffix used in their gene name, and their numerosity in the human genome. Adapted from (Seal et al., 2020).

| LncRNA subtypes | Description | Nomenclature characteristic | Abundance in human genome |
| --- | --- | --- | --- |
| Novel transcripts | Newly identified lncRNAs | ENSG | 14,320 |
| Name assigned lncRNAs | Already studied lncRNAs, that have been attributed an official and coded gene name | none | 1,040 |
| Antisense lncRNAs | lncGs that totally or partially overlap the genomic sequence of a protein-coding gene on the opposite strand | -AS1<br>-AS2<br>... | 1,848 |
| Divergent Transcripts | lncGs transcribed from a bidirectional promoter in the opposite direction to a protein-coding gene | -DT | 600 |
| Intergenic lncRNAs | lncGs not overlapping a protein-coding gene, not sharing a bidirectional promoter with a protein-coding gene, and not hosting microRNA or snoRNA genes | LINC | 2,274 |
| Intronic Transcripts | lncGs transcribed entirely from an intron of a protein-coding gene on the same strand | -IT1<br>-IT2<br>... | 127 |
| MicroRNA Hosting Genes | lncGs hosting microRNA genes | -HG1<br>-HG2<br>... | 86 |
| Overlapping Transcripts | lncGs overlapping protein-coding genes on the same strand | -OT1<br>-OT2<br>... | 15 |

A substantial portion of lncGs (approximately 70.5%) is categorized as “novel transcripts” in line with the recognition that numerous lncRNAs remain unexplored and uncharacterized, thus lacking an official gene name or functional classification. While being the most prevalent lncRNA class in almost all chromosomes, with only a few exceptions (**Supplementary Figure 1c**) novel transcripts tend to be relatively shorter, with their gene having a mean length of 29,690.81 nt (**Supplementary Figure 1d**).

The “Long intergenic” (LINC) class of lncRNAs makes up approximately 11.2% of all lncGs and is associated with a lower GC content compared to other lncRNA subtypes, particularly evident in their 3’ end (**Figure 2d**).

The category “Overlapping transcripts” includes lncRNAs predominantly located on chromosome 3 (**Supplementary Figure 1c**). A notable characteristic of this group is the variable length of genes with a mean of 61,152.1333 nt (SEM= 31,091.165). Additionally, these transcripts exhibit a relatively constant GC content along their sequence, ranging from a maximum of 54% to a minimum of 48%, which could relate to their intrinsic characteristic of overlapping a PCGs (**Figure 2d**).

MicroRNA hosting genes are predominantly located on chromosomes 21 (24%) and 20 (15.1%), while they are absent on chromosomes 3 and Y. These genes are also the longest ones among lncGs, with an average length of 118,660.023 nt, and their transcripts display the lowest GC content at the 3’ end compared to all other lncRNAs (33.6%) (**Figure 2d**).

Finally, “divergent transcripts” represent the lncRNA class with the most pronounced difference in GC content between their extremities, ranging from 60.7% at the 5’ end to 36.3% at the 3’ end (**Figure 2d**). Since these transcripts are transcribed in the opposite direction to PCGs, it remains to be determined whether the skewed GC content is functionally related to their bidirectionality.

It is worth noting that this classification is not exhaustive, and it is influenced by several factors, including the diverse and heterogeneous features considered, the substantial portion of unannotated transcripts, and the potential misclassification of lncRNAs that have already been characterized (name-assigned lncRNAs). Nevertheless, it could serve as a step towards achieving a clearer functional distinction between lncGs based on their genomic, structural and/or functional characteristics.

### Regulatory and functional elements of lncGs

Most of the lncRNAs are cleaved and polyadenylated at their 3’ends, like the majority of the RNA polymerase II-transcribed genes. Others, instead, are stabilized by alternative mechanisms, including RNase P cleavage (Brown et al., 2012; Wilusz et al., 2012), or processed on both ends by the snoRNA machinery (Yin et al., 2012). Upon analyzing the presence of canonical polyadenylation (polyA) sites (CA) and polyA signals (AATAAA) within the genomic sequences of lncGs, we discovered that nearly 28% (5,587) of lncGs exhibit the presence of either a polyA site or a polyA signal. This relatively low percentage aligns with other reports indicating that lncRNAs are predominantly non-polyadenylated (Schlackow et al., 2017). The mean distance between the polyA site and the polyA signal was 20.7 nt and 20.3 nt for protein-coding and lncG sequences respectively (**Supplementary figure 2b**), indicating no evident differences between the two classes in terms of the organization of polyadenylation elements within their genomic sequences.

As polyA regulates mRNA export to cytoplasm and translation, we analyzed the presence of canonical polyA sites and signals in lncGs in relation to the subcellular localization of their related transcripts. By querying the “RNAlocate” tool (http://www.rna-society.org/rnalocate/) database, which includes 589 annotated lncRNAs along with their subcellular localization, we found no significant correlation between the presence/absence of polyA signals and the localization of specific lncRNAs into nuclear or cytoplasmic compartments (Fisher exact test p-value=0.126).

Similar to PCGs, lncGs are composed of one or more exons and introns within their genomic sequence. Some lncGs can be transcribed into different isoforms due to alternative splicing, variations in 5’ or 3’-ends, exon number, or the presence of introns that are not removed by the splicing machinery. Most PCGs are polyexonic, with only 4.5% being monoexonic. On average, they consist of 70 exons (median=34) resulting in a total of 171,410 distinct annotated transcripts (**Table 2**). Each gene encodes an average of 8.5 diverse RNA isoforms (median=6), produced by alternative splicing. We found RBFOX1 (RNA Binding Fox-1 Homolog 1) to be the longest PCG (2,473,539 nt), coding for 40 different splicing variants.

**Table 2.** Genomic elements of protein-coding and non-coding genes, including transcript isoforms and exon number.

|  | Protein-coding<br>genes/transcripts | Long non-coding<br>genes/transcripts |
| --- | --- | --- |
| N° of genes | 20,065 | 20,310 |
| N° of transcripts | 171,411 | 59,928 |
| Mean exons per gene | 70.1 | 11.6 |
| Median exons per gene | 34 | 3 |
| N° of mono-exonic genes | 973 | 3,173 |
| Mean exons per transcript | 8.2 | 3.8 |
| Median exons per transcript | 6 | 3 |
| Mean transcripts per gene | 8.5 | 3.1 |
| Median transcripts per gene | 6 | 1 |

From the lncRNA perspective, the longest gene is annotated as NRXN1-DT, a divergent transcript with a genomic length of 1,375,317 nt. This lncG is located on the positive strand of chromosome 2 and is transcribed in its full length into a 3,939 nt monoexonic RNA. Considering all lncGs, 15.6% of them are monoexonic while the others exhibit a varying number of exons, with a mean number of exons in the encoded transcripts that is lower than that of protein-coding transcripts (3.8 exons for lncRNAs, 8.2 for protein-coding transcripts). The lncRNA with the highest number of exons is SNHG14, a spliced paternally imprinted lncRNA located in the Prader-Willy critical region, comprising 59 different exon combinations and contributing to the transcription of 142 different splicing variants. However, the lncRNA producing the highest number of splicing variants is the antisense PCBP1-AS1 (the second in rank for number of exons) with 296 annotated transcripts. In total, considering all lncGs, they possess a mean of 3.1 distinct splicing variants, contributing to a total of 59,928 long non-coding transcripts.

Taken together, these results underscore potentially crucial functional features of lncRNAs that distinguish them from PCGs (**Table 2**). These features include polyA, as well as a lower number of exons within their sequence, and a reduced number of splicing variants.

### Nucleotide variations in long non-coding RNA genomic sequences

Nucleotide variation can significantly impact gene functionality (Haraksingh and Snyder, 2013; Karczewski et al., 2020), including lncGs (Esposito et al., 2023). Mutations occurring in promoter regions or splicing sites could profoundly affect their expression, while mutations affecting binding regions could impact the ability of the gene product to interact with RNAs, DNA sequences, and proteins.

Our analysis of the human single nucleotide variations (SNVs) cataloged in dbSNP (Sherry et al., 2001), identified approximately 130 million single nucleotide polymorphisms (SNPs) associated with lncG regions. Among clinically annotated single nucleotide substitutions, transversions were the most common, accounting for 63.3% of all SNVs in lncGs and 63.7% in protein-coding genes (PCGs) (**Figure 3 a**). We observed no significant differences in the frequency of SNVs between lncGs and PCGs.

**Figure 3.**
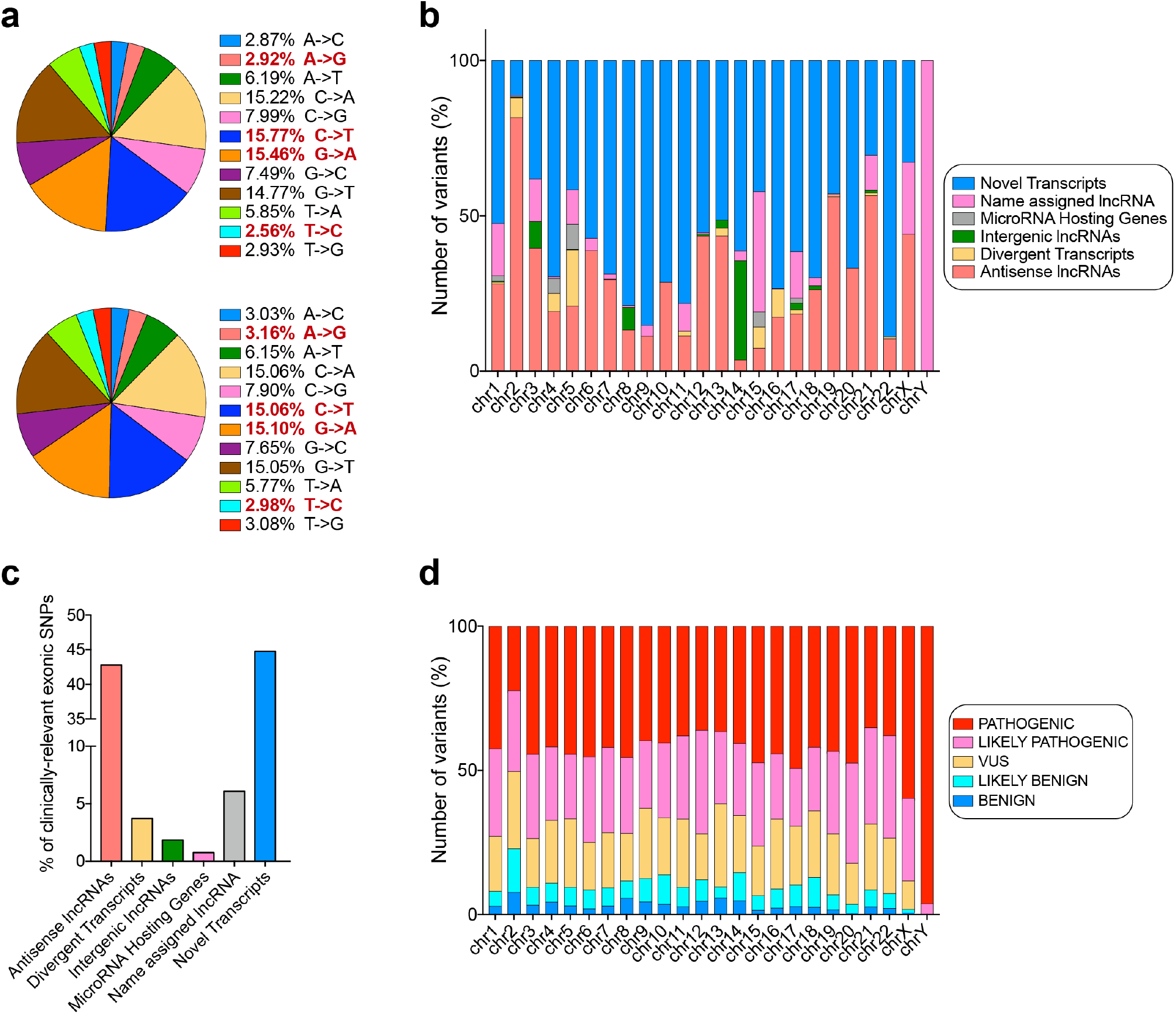
Analysis of SNPs across long non-coding genes. **(a)** Piechart showing the percentages of transitions and transversions in lncGs (upper piechart) and PCGs (lower piechart). Transitions are bolded and red colored. **(b)** Percentages of variants on the distinct chromosomes divided by long non-coding subtype. **(c)** Barplot showing the percentage of clinically-associated SNPs in the distinct long non-coding RNA subtypes. **(d)** Clinically-associated variants (%) in lncGs, grouped by pathogenicity for each chromosome.

According to Ensemble annotations, 14,303 variants in lncGs are classified as clinically relevant, with 36.4% labeled as pathogenic, 27.4% as likely pathogenic, and 22.1% as variants of uncertain significance (VUS). We then analyzed lncRNA subtypes to determine those more susceptible to clinically associated mutations (**Figure 3 b**). Intronic and overlapping transcripts showed no clinically associated variants. However, two categories stood out for their susceptibility to SNPs: antisense RNAs and novel transcripts. Antisense transcripts were highly enriched on chromosome 2, comprising the majority of variants observed in this region. By contrast, the Y chromosome exhibited only variants associated with name-assigned lncRNAs.

Notably, divergent transcripts were predominantly affected on chromosome 2, accounting for 56% of total mutations in this RNA class. Conversely, variants in long intergenic non-coding RNAs (LINCs) were primarily distributed across chromosomes 3, 8, and 14, even though LINCs are predominantly located on chromosome 13. MicroRNA-hosting genes appeared to be the least affected by SNPs, constituting just 0.74% of all SNPs, with most of these variants located on chromosomes 1, 4, 5, 15, and 17 (**Figure 3 c**). Overall, only 0.01% of SNPs in lncRNAs are classified as clinically relevant.

When examining the pathogenicity of SNVs across individual chromosomes, similar patterns emerged (**Figure 3 d**). The Y chromosome stood out, containing 26 out of 27 pathogenic variants, along with one likely pathogenic variant. Chromosome 2 had the lowest percentage of pathogenic variants. Following the Y chromosome, the X chromosome exhibited one of the highest proportions of pathogenic variants (59.7%). This chromosome is of particular significance due to the presence of XIST, a well-characterized lncRNA responsible for X-chromosome inactivation in female mammals (Markaki et al., 2021). Interestingly, while chromosome 2 had the highest absolute number of clinically associated variants (n=4,552), these represent 31.8% of all variants in lncGs. Altogether, these findings suggest that lncRNA genomic regions are characterized by an uneven distribution of SNPs, with clinically significant variants being concentrated on distinct chromosomes, each exhibiting different levels of pathogenicity.

## Discussion and conclusions

Studying lncGs is a challenging task as their classification substantially relies on general features associated with multiple parameters, including length, position relative to coding genes, structural properties and activity. Compared to mRNAs, where the genetic code guides functional inferences, lncRNAs are more puzzling to characterize due to their complex transcriptome and processing, lack of evolutionary constraints, and limited structural/functional correlations. Therefore, precise and thorough annotation is crucial for fully exploiting the potential of noncoding genomic data and achieving significant advancements in biomedical research. These issues require experimental approaches for detailed lncRNA annotations, highlighting the importance of their genomic architectural definitions as foundational references.

To advance our understanding of lncRNAs, we sought to dissect their genomic organization and correlate it with existing annotation and functional elements. Our objective was to delineate the current genomic landscape of lncGs, effectively creating a virtual long non-coding RNA karyotype.

By capturing and processing data from the available human gene set in GENCODE (see **Methods**), we show a screenshot of the current numerology of lncGs. Their abundance is comparable to that of protein-coding genes (PCGs), with around 20,000 hits and an average length of about 31,600 nt, which is consistently shorter than PCGs (72,400 nt).

Moving to transcripts and considering the effect of alternative splicing on the RNA output, a total number of 59,928 long non-coding RNA variants was calculated. Transcript length is one of the most crucial features for distinguishing lncRNAs from other non-coding species, and it has been originally employed for their identification. However, the use of this parameter revealed a heterogeneous population of molecules, spanning from a few hundreds to hundreds of thousands of nt in length. Overall, their lengths appear to be related to chromosome size, yet discrepancies exist, particularly in chromosomes 21 and Y, where lengths are longer than expected, and in chromosome 1, with shorter lengths than expected. Whether the size of lncGs is a regulative determinant for the functionality of the lncRNA mature form is a plausible factor which needs further exploration, as a significant portion of the genetic material was predicted to be spliced out after transcription.

From a topological standpoint, lncGs are generally evenly distributed across the genome, albeit with some exceptions observed on specific chromosomes and localized density peaks on distinct chromosomes. Notably, chromosome 2 was found to harbor the highest density of lncGs and is more affected by clinically relevant SNPs, although they appear to be less pathogenic compared to those on other chromosomes. Additionally, we identified positional clustering of lncGs, particularly on chromosomes 21 and Y. This observation may suggest the existence of genomic loci dedicated to lncRNAs, which could be further and specifically focused through quantitative gene expression analyses to assess whether this positional clustering underlies functional implications. Indeed, a few reports explored the role of intronic or intergenic lncG clusters co-expressed and functionally synergistic in brain development (Wang et al., 2019) and cancer settings (Tomita et al., 2015; Zhou et al., 2021), respectively. This indicates clustering as an opportunity for lncRNA coordinated expression, combinatorial activity and clinical significance in physiological and diseased mechanisms. However, the biological functions of most lncRNA clusters remain largely unclear (Sun et al., 2013).

Making a comparison with coding transcripts, we also inspected some functional elements within lncG sequences that could potentially affect their transcription. These elements, include a distinct pattern of GC content, that was particularly evident when comparing the 3’ and 5’ ends of these genes. Finally, we examined the number and quality of nucleotide variations affecting lncRNA sequences. While human genetics studies predominantly focus on coding genes, often associated with distinct genetic or multifactorial diseases, we identified a number of clinically relevant SNPs impacting lncGs. It is important to note that some of these variations could affect antisense coding genes or regulatory regions relevant to coding genes. However, as the expression and function of lncRNAs could also be influenced by the presence of these SNPs, it is conceivable that lncRNAs may play a role in certain diseases. In this context, the study of expression quantitative trait loci (eQTL), such as that conducted by the FANTOM project (Hon et al., 2017; Yip et al., 2022), represents an invaluable source of information for understanding the implication of lncRNA nucleotide variations in genetically determined conditions and phenotypes.

## Methods

### Datasets used in this study

Annotation files related to both human lncRNA (Long non-coding RNA gene annotation) and coding (Comprehensive gene annotation) genes were downloaded from GENCODE Release 46 (GRCh38.p14) in GTF format. Sequence file of long non-coding transcripts (Transcript sequences) was downloaded from GENCODE in FASTA format. Polyadenylation data of all human transcripts (PolyA feature annotation) was downloaded from GENCODE in GTF format. SNPs data was downloaded from Ensembl release-110 in VCF v4.1 format.

### Data processing

Analyses have been performed in the R environment. GTF files were read and processed using *rtracklayer* package (Lawrence et al., 2009). FASTA files were read and processed using *Biostrings* (Pagès et al., 2021). Other R packages used for the analysis are *biomaRt* for gene and transcripts ID conversion, *VariantAnnotation, GenomicRanges* and *seqinr* for sequence analyses (including GC content computation by sliding windows and SNPs analysis).

Long non-coding RNA and protein-coding genes distributions (peaks) were computed with a 1 Megabase window and reported as rainfall plots using the *circlize* library. In peaks quantification, these are the regions where the gene density exceeds a certain threshold, indicating that they are areas of unusually high concentration of either lncGs or PCGs. The threshold was set at the 95th percentile of the gene density distribution (*quantile(density_lnc$density, 0*.*95)* for lncGs and *quantile(density_cod$density, 0*.*95)* for PCGs). This means that the top 5% of density values are considered “TRUE peaks” and represented as triangles, while “FALSE peaks” were represented as circles.

Genomic elements (polyadenylation and exons) were retrieved from the annotation files and processed accordingly to compute the statistics (filtering in all polyadenylated genes or exon— annotated genes respectively).

GC content was calculated by summing the number of G and C nucleotides for each transcript sequence from FASTA files. GC skew computation was performed by implementing a function that calculates the GC content on a window size of 10 nucleotides. The results are divided by 20 chunks for each transcript sequence and then the average GC content is calculated for each chunk.

LncRNA subclassification was performed on each annotated gene name, as follows: gene names starting as “LINC” were classified as “Long Intergenic lncRNA”; gene names starting with “ENSG” were annotated as “Novel Transcripts”; gene names ending with “-HG”, “-AS^*^”, “-DT^*^”, “-IT^*^”, “-OT^*^” were annotated as “MicroRNA Hosting Genes”, “Antisense lncRNAs”, “Divergent Transcripts”, “Intronic Transcripts” and “Overlapping Transcripts”, respectively. All the other genes were annotated as “Known lncRNAs”.

The analysis of the distance between polyadenylation sites and signals was performed on the genomic coordinates of each polyadenylated gene as annotated in the genomic files (start sites and end sites). For each chromosome, SNP analysis was performed on SNP data from Ensembl Variation filtered for SNPs falling on genomic regions covered by lncG or PCG sequences. Clinically-associated SNPs were retrieved as VCF files from Ensembl.

Code is available by the authors upon reasonable request.

## Declarations

### Availability of data and materials

The datasets generated and/or analyzed during the current study are available in the following repositories:

GTF file with lncRNA annotation, Sequence file in FASTA format of long non-coding transcripts, and Polyadenylation data of all human transcripts: GENCODE Release 46 (GRCh38.p14) (https://www.gencodegenes.org/human/release_46.html); SNPs data in VCF v4.1 format: Ensembl release-110 (http://www.ensembl.org/Homo_sapiens/Info/Index); Code used during the current study is available from the corresponding author on reasonable request.

## Authors’ contributions

CRediT author statement: Conceptualization AP, MB. Methodology, Software, Validation, Data Curation, and Visualization: AP. Formal analysis, Investigation and Resources: AP, MB. Writing - Original Draft: AP. Writing - Review & Editing: AP, GB, PL, MB. Supervision and Project administration: AP, PL, MB. Funding acquisition: PL, MB.

